# Compaction and swelling of single stretched DNAs driven by molecular crowding

**DOI:** 10.64898/2025.12.09.693115

**Authors:** Paritosh Gupta, John F. Marko, Vittore F. Scolari

## Abstract

We present a theory for the effects of osmotic pressure exerted by macromolecular crowders, on a double-stranded DNA or other semiflexible polymer extended by tension. Our results predict the force and crowder-density dependence of the polymer extension. The lowest-order effect is a crowder-dependent compression that counteracts the external stretching force, which can collapse the polymer at a crowder-dependent critical force *f*^*^. This compression is dependent on crowder radius *r* and density *ϕ*, with higher densities and smaller radii having greater effects on the force-extension curve. At first order in perturbation theory we also find a fluctuation-dependent correction to the depletion volume, which can overwhelm the simple compression effect for large crowders, leading to expansion of the polymer.

---

The genetic material of living cells is confined in compartments where it shares its space with a large number of other macromolecules fundamental for cellular functions. Many of these macromolecules, such as transcription factors, DNA and RNA polymerases, nucleosomes, specific enzymes and molecular motors associate with DNA, forming nucleoprotein complexes along *chromatin* in eukaryotes, or the *nucleoid* in bacteria. In both these cases, one can consider the chromosome fiber as a polymer: in eukaryotes, chromatin is at base a string of nucleosomes, while in bacteria the nucleoid fiber consists of DNA decorated by DNA-binding proteins. However, many other macromolecules near the genome do not bind to the nucleoid or chromatin fibers, but do share the same constrained space inside the cell, and interact with the genome via steric exclusion.

These “free molecules”, that we call *crowders*, are characterized by a high diffusive mobility, and produce osmotic pressure. This pressure is exerted towards the exterior of the cell, on the plasma or cell membrane, and is a key factor in the regulation of overall cellular volume [1]. The same pressure is applied towards the interior, and ultimately on the genetic material. Analogous to how depletion forces by macromolecules can induce attractive interactions between larger colloidal particles [2], these osmotic depletion forces can compact chromatin and induce its phase separation [3, 4].

Experiments *in vivo* have observed osmotic shock-driven condensation of the nucleoid in *Escherichia coli*, in a reduction of nucleiod volume following an increase of salt concentration in the extracellular medium [5]. The osmotic shock drives water out of the cell and causes a direct reduction in cell volume. In turn, this induces an increase in concentrations of the crowders in the cytoplasm. It was further shown that the volume reduction is considerably smaller for the cytoplasm than for the nucleoid, hinting at a cooperative effect which can be thought about as a type of phase separation (i.e., via considering the energetic cost of expanding the nucleoid volume).

Complementary *in vitro* experiments using purified bacterial nucleoids and a set of crowders solutions covering a large range of densities and sizes have shown that crowders at physiological levels can directly compact the nucleoid [6–8], indicating that crowding can be a key factor in chromosome condensation. These observations indicate that crowders at physiological densities can regulate the level of condensation of the bacterial chromosome fiber into the nucleoid, allowing its membraneless compartmentalization.

Similar phenomena have been observed for eukaryotic chromatin in human cell lines: an increase in density of molecular crowders has been correlated to chromosomal mitotic condensation [9] following the dissolution of the nuclear envelope. Osmotic shock has been shown to condense mitotic chromosomes similar to how it affects the nucleoid in *E. coli* [9]. Further, it is possible that the effect of crowding-induced liquid-liquid phase separation of key chromatin-associated proteins [10] is relevant for the chromosome’s three-dimensional compartmentalization in silent and transcribed regions.

Experimental results and theory have been complemented by molecular dynamics simulations reproducing crowders mediated compaction of polymer models *in silico* [5, 11–14]. Typical simulations of this type consist of *L* spherical beads of radius *r*_0_ connected by springs of a similar extension, immersed in in a bath of small spherical crowders of radius *r* with density *c*. These simulations identified that the crowder volume fraction *ϕ* = 4/3 *πr*^3^*c* as well as the relative crowder size *r*/*r*_0_ are fundamental in setting the strength of the crowder-induced compacting force. Crowding-driven collapse of unstretched semiflexible polymers has also been numerically observed and quantified [15, 16]. While simulations has demonstrated that crowders have the ability to compact and collapse polymers, there are limits of this approach towards a quantitative theory. In general, molecular dynamics simulation results, as all empirical derivations, are valid only on the range of parameters that were investigated, and this methodology does not discern the theoretical determinants of the range of validity. Furthermore, it is a challenge to improve this type of simulation, since a reduc-tion of *r*/*r*_0_ by a factor *x* at fixed *ϕ* entails an increase of the number of simulated particles of 𝒪 (*x*^3^). Similarly, an increase of the polymer length by a factor *x* entails, due to scaling of polymer dynamics, an increase of the simulated volume and number of simulated particles of 𝒪 (*x*^5/2^) . On a more fundamental level, the bead-spring model is not a good polymer to simulate DNA (or DNA-based chromosome fibers), as DNA is a significantly slender polymer with bending stiffness (persistence length) much longer than its cross section, and this makes quantitative matching of parameters in published molecular dynamics simulations to experiments problematic.

One can carry out molecular pulling experiments with high precision *in vitro*. These types of experiments can be used as a tool for quantitative study of crowding agents, by quantifying how the crowders change the force-extension behavior of DNA, or in principle isolated chromosome fibers. As an example, a recent experiment has shown how crowding affects DNA supercoiling, suppressing its tendency to strand-separate under negative torsional stress [17].

Here we present a theory of effects of molecular crowding on the force-extension relation for a semiflexible polymer or worm-like chain (WLC), in the strong stretching regime, valid for large enough forces that the polymer is extended, and low enough crowder density so that an ideal crowder gas approximation for osmotic pressure is also reasonable. We will include quantitative discussion for the key experimental case of double-stranded DNA.

We consider a worm like chain (WLC, equivalently a semiflexible polymer) stretched by external tension *f*, described by its tangent vector 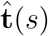 over the arc-length coordinate *s* spanning the full length *L* of the chain. In coordinates with cylindrical symmetry around the 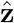 pulling direction, we write the energy as the sum of the bending energy of the WLC with bare persistence length *A*, the usual coupling of the externally imposed pulling force to the molecule extension 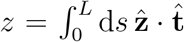, and an osmotic term due to the crowders in the solvent. This osmotic contribution is simply the product of the crowder concentration *c* and the volume made sterically inaccessible to the crowders by the presence of the polymer chain *δV* :

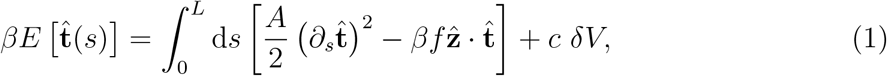

for more details on the derivation and underlying assumptions see the Appendix.

In the zero-crowder-concentration limit (*c* → 0 or equivalently, volume fraction *ϕ* = *vc* → 0) this becomes the standard WLC model under pulling force [18]. The model has a characteristic force *f*_*c*_ = *k*_*B*_*T/A* that corresponds to the force at which the Pincus blob radius [19] is equal to the persistence length of the chain. The force *f*_*c*_ corresponds to WLC extension of roughly 50% of its maximum. For pulling forces *f* > *f*_*c*_, an expansion in inverse powers of *f* gives *z* as the full chain extension *L*, minus a force-dependent term that is proportional to the intensity of the chain fluctuations perpendicular to the pulling force ⟨*z*⟩ /*L* = 1 − (4*f* / *f*_*c*_)^−½^ + · · · as depicted in Fig. 1A, black line.

**FIG. 1.**
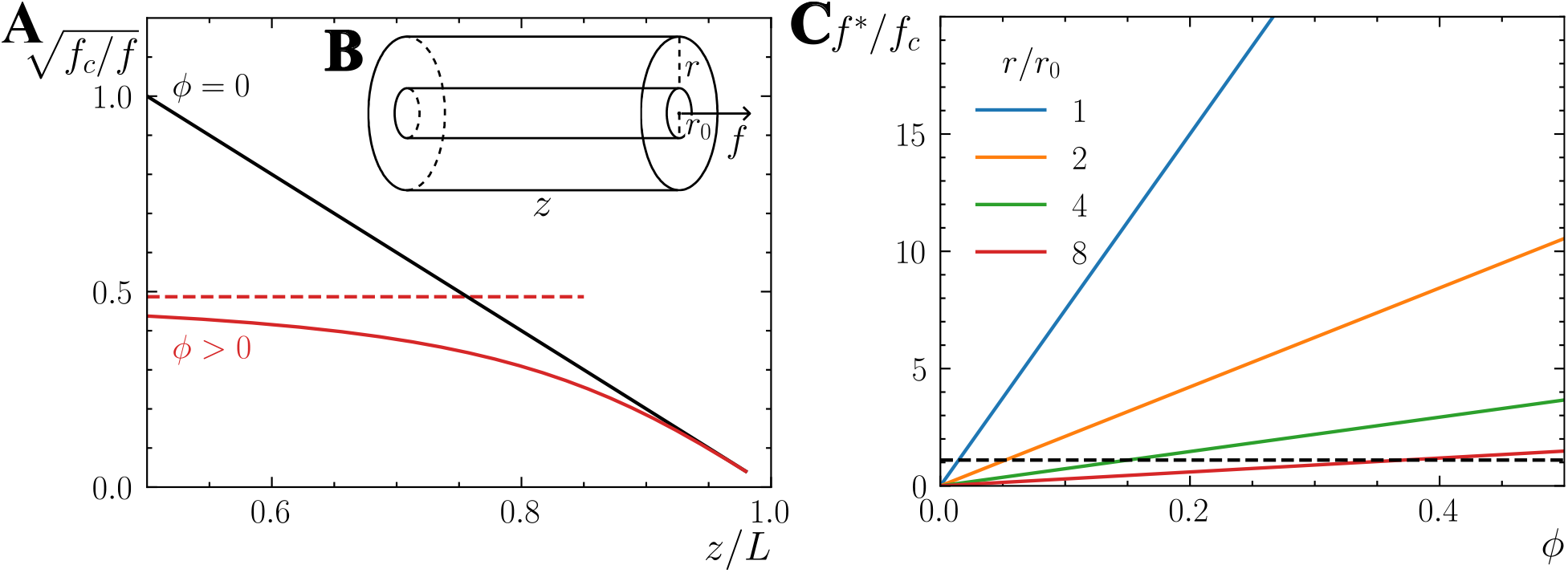
Effective force in the rigid cylinder approximation. **(A)** Force extension relationship, without crowders (*ϕ* = 0) *vs* with crowders (*r/r*_0_ = 2 and *ϕ* = 0.2). The crowders reduce the effective stretching force by a factor *f*^∗^, indicated by the dashed red line. **(B)** Rigid cylinder approximation for the stretched WLC. Neglecting fluctuations, the volume depleted by the chain to crowders is a cylinder of extension *z* and radius equal to the sum of the cross-radius of the polymer *r*_0_ and the crowder’s radius *r*. **(C)** Collapse force *f*^∗^ as function of crowder density *ϕ*, for varied crowder size *r*. Smaller crowders have a stronger collapsing effect. Values of *f*^∗^*/f*_*c*_ = 1.1 are indicated by dashed black line; below this the collapsing effect of crowders on the force-extension relationship is small and difficult to detect.

To calculate *δV* in Eq. 1, we consider first the approximation where we neglect the effects of the transverse fluctuations of the chain, taking it to be a rigid rod of radius *r*_0_ and variable elongation *z*. In this configuration, the volume *δV*_*cyl*_ inaccessible to the crowders is a cylinder of radius *R* = *r*_0_ + *r* and of the same length as the polymer extension, as depicted in Fig. 1B:

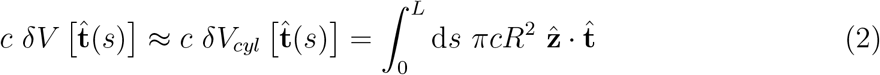

Here the crowders are considered to be spheres of radius *r*, but the theory can be generalized to other shapes of crowding molecules. This is the zeroth-order starting point for dealing with the transverse fluctuations, which modify the volume excluded to the crowders.

The osmotic force acting on the semiflexible chain as a whole tends to reduce its extension *z* by acting against the effect of the external force *f* with a term that has exactly the same dependence on the conformation 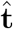 as the pulling term (see Eq. 1). This corresponds to an additive reduction of the effective pulling force by a crowder-dependent *f*^∗^, as depicted in Fig. 1A, red line, with

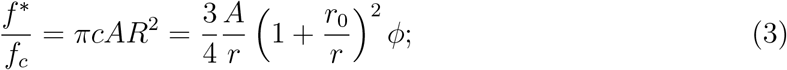

the dashed red line in Fig. 1A indicates the value of the ordinate corresponding to *f*^∗^. All crowding-dependent terms in this rigid cylinder approximation are in this effective force term 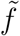:

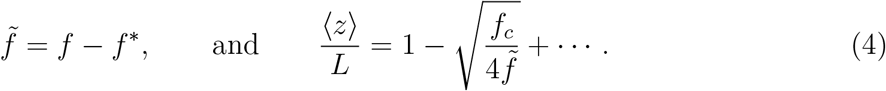

This result predicts a chain collapse instability for external forces *f* < *f*^∗^, where the net effective pulling force goes to zero. The larger the critical force, the larger is the effect of crowders on the force-extension relationship. The effects are proportional to the crowder volume fraction *ϕ*, and become more significant for smaller crowder sizes *r/r*_0_, as depicted in Fig. 1C. While we expect the chain to collapse at effective forces 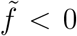, once collapse occurs, we expect higher-order fluctuation terms, effects of polymer looping, surface tension and nematic effects to become important in defining the condensed “phase”. These effects are not accounted for by our theory, since its domain of validity is 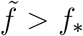 (and *f*_*c*_); in this regime the polymer is extended so these nonlinear effects do not play a role.

We can estimate effects of crowders for the specific case of double-stranded DNA in physiological salt conditions, considering that the persistence length is *A* = 50 nm, giving characteristic force *f*_*c*_ ≃ 0.08 pN, and that the radius of DNA is *r*_0_ ≃ 2 nm, for crowders of radius *r* = 16 nm it takes a volume density of at around *ϕ* > 0.4 to observe relative effects (*f*^∗^ − *f*_*c*_)*/f*_*c*_ above 10% (crossing of red and dashed black lines in Fig. 1C), and it takes larger volume densities to see effects at higher forces. For substantially larger crowders, the jamming of the solvent, significant at *ϕ* > 0.5, would make it very difficult to see any polymer collapse at any values of concentration and pulling forces. On the other hand, smaller crowders have an observable effect at lower densities, *i*.*e*. for *r* = 3 nm, we observe a 10% effect already at *ϕ* > 0.06. This size of crowders is comparable in size to the radius of proteins and protein complexes in the nucleoplasm: therefore we expect crowding-induced collapse to be relevant *in vivo* where protein concentrations are in the range of *ϕ* > 0.1.

We now add the effect of WLC bending fluctuations that were neglected in the previous computation. The polymer, pulled by an effective force 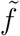 in the 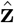 direction, has fluctuations in the 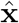 and **ŷ** directions orthogonal to the force, as depicted in Fig. 2A. The orthogonal displacement of the chain **u**_⊥_(*s*), defined by the orthogonal component of the tangent vector **t**_⊥_(*s*) through *∂*_*s*_**u**_⊥_(*s*) = **t**_⊥_ has fluctuations with a force-dependent correlation length 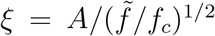 [18], or explicitly ⟨**t**_⊥_(*s*_0_) · **t**_⊥_(*s*)⟩ = (*ξ/A*) exp[−(*s* − *s*_0_)/*ξ*] relative to any point *s*_0_ along the polymer. Consider a crowder touching the chain at some particular position *s*_0_: the trajectory of the chain **u**_⊥_(*s*) will have a rigid-rod conformation in the direction **t**_⊥_(*s*_0_) for |*s* − *s*_0_| < *ξ*, and will then begin to random walk in the transverse directions for larger values of |*s* − *s*_0_|. This configuration will be excluded if this trajectory “overlaps” with the same crowder at a second point away from *s*_0_.

**FIG. 2.**
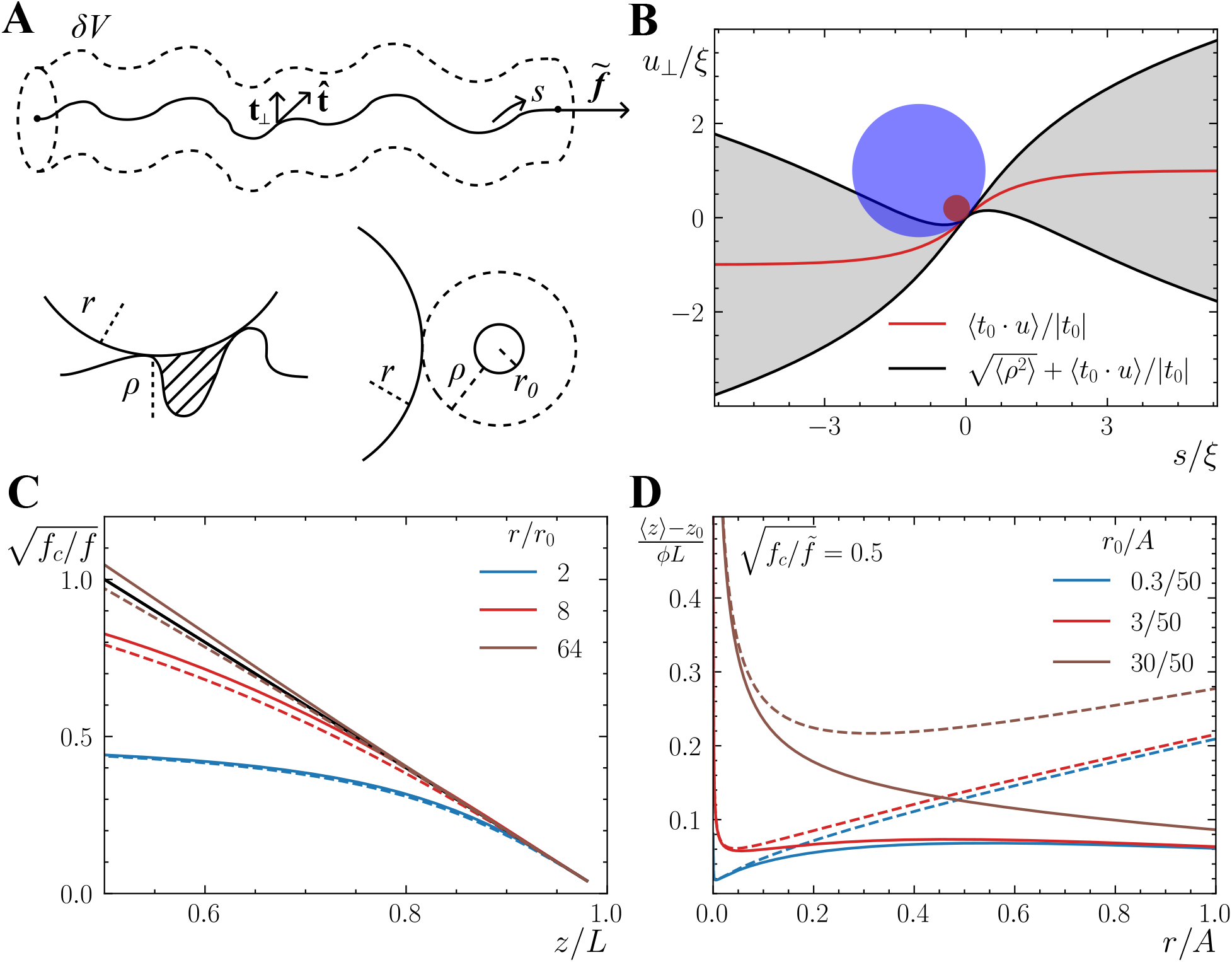
Effects of chain fluctuations. **(A)** The polymer fluctuates perpendicular to the pulling force, generating additional excluded volume *δV* not accounted for by the rigid cylinder approximation. We account for this by an additional fluctuation dependent radius *ρ*. **(B)** 1D illustration of calculation of ⟨*ρ*^2^⟩ at fixed *r/r*_0_ and *ϕ*. Two spherical crowders of different radius touch the polymer at *s* = 0. The non-zero mean component of the WLC trajectory along the **t**_⊥_(0) direction does not contribute to depletion while standard deviation does. **(C)** Force extension relationship, as in Fig. 1A (*ϕ* = 0.2 and *r/r*_0_ = 2). Dashed line shows zeroth-order contribution of crowders; solid lines accounts for fluctuation-dependent corrections. **(D)**. Polymer fluctuation-dependent contribution to the force-extension relationship as function of *r/A* at fixed effective force 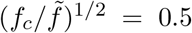, from Eq. 10 (solid lines, red line corresponding to DNA *r*_0_ value). Dashed lines show *λ* ≪ 1 approximation (Eq. 11).

To account for this we consider this effect as an increase of the radius of the cylinder of depletion by an additional fluctuation dependent distance *ρ*, representing the additional volume inaccessible to the crowders as depicted in Fig. 2A. The depletion volume is

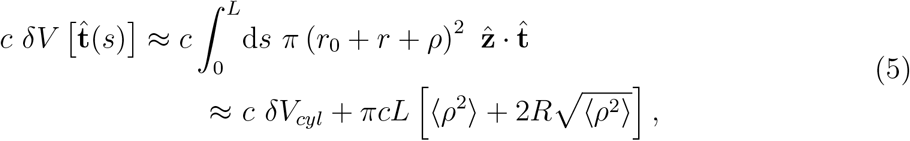

where we have used a perturbation expansion of the partition function to first order in crowder volume fraction *ϕ*.

To calculate ⟨*ρ*^2^⟩ we consider the orthogonal displacement **u**_⊥_(*s*) of the WLC conformation over distances comparable to the crowder size *r*, which should be compared to the correlation length for bending and displacement fluctuations *ξ*. For small crowders *r* < *ξ* care must be taken: the tangent vector has strongly correlated tilt over distances of the crowder size, expressed by a nonzero value of ⟨**t**_⊥_(0) · **u**_⊥_(*s*)⟩/⟨|**t**_⊥_(0)|^2^⟩^1/2^. This quantity is the component of orthogonal displacement in the direction of the tangent vector tilt, **t**_⊥_(0)/ |**t**_⊥_(0)|, and can be computed to be

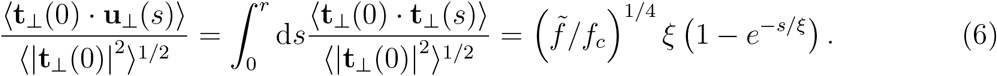

Subtracting this from **u**_⊥_(*s*) yields displacement fluctuations in the tilted reference frame, correctly expressing the area transverse to the polymer that is excluded to the crowders (Fig. 2B).

The fluctuation-corrected effective radius valid for all *r* follows as

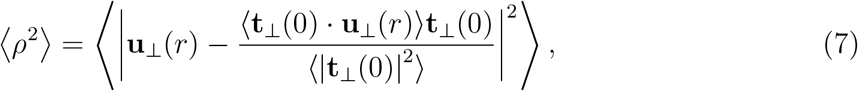

which can be written in terms of a dimensionless scaling function *g*(*λ*) of scaled crowder size *λ* = *r/ξ*:

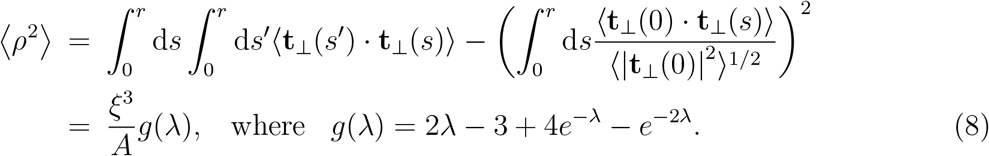

The exclusion radius depends on fluctuations and is force-dependent; for sufficiently large force and large crowders (*r > ξ* or *λ* > 1), *g*(*λ*) = 2*λ* +· · ·, with asymptotic behavior ⟨*ρ*^2^⟩ = 2*k*_*B*_*Trξ*^2^*/A*+· · ·, compatible with the usual WLC transverse size result [18]. In the opposite small-crowder limit (*r* < *ξ* or *λ <* 1), *g*(*λ*) = 2*λ*^3^/3 + · · · one has ⟨*ρ*^2^⟩ = 2*r*^3^/(3*A*) + · · · . In this latter *λ* → 0 limit, the force-dependence disappears, compatible with the analysis of transverse fluctuations of a WLC confined to a narrow tube [20] (note that without removal of tangent tilt one has the incorrect small-*λ* behavior *g*(*λ*) ≈ *λ*^2^ and ⟨*ρ*^2^⟩ ≈ *r*^2^*ξ/A*, resulting in unphysical force dependence persisting as *r* → 0, *i*.*e*., as the crowders vanish).

The partition function for the WLC including crowders can now be written down, shifting 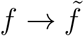 to account for *δV*_*cyl*_, and adding the excluded volume from (5) including (8):

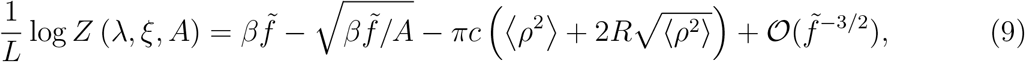

The force-extension relationship follows as 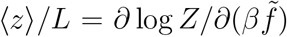, leading to correction of the zeroth-order force extension relationship of Eq. 4 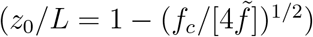:

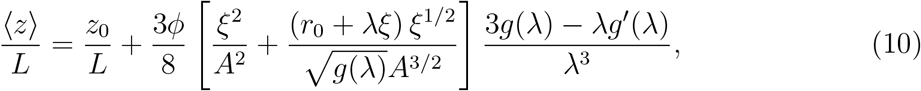

where ⟨*z*⟩*/L* is a function of the force-dependent bending fluctuation correlation length *ξ*, the volume fraction of the crowders *ϕ*, their size via *λ* = *r/ξ*, the polymer stiffness (persistence length) *A*, and the bare polymer radius *r*_0_. This result is plotted in Fig. 2C.

The fluctuation correction to ⟨*z*⟩*/L* is always positive, since 3*g*(*λ*) − *λg*^*′*^(*λ*) is positive for all *λ*. We define a characteristic force *f*_*λ*_ = *f*_*c*_*A*^2^*/r*^2^ at which *λ* = 1, which is the boundary between the two scaling regimes for the fluctuation corrections. For 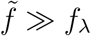, both *g*(*λ*) as well as 3*g*(*λ*) − *λg*^*′*^(*λ*) vary ∼ *λ*. In this regime, the fluctuation correction goes to zero as 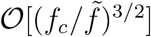 asymptotically, as such the polymer’s extension always attains its contour length for infinite pulling force. It also monotonically decreases toward zero for large crowders as 𝒪 [(*A/r*)^3/2^] as is physically reasonable.

In the opposite limit 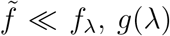 varies ∼ 2*λ*^3^/3 while 3*g*(*λ*) − *λg*^*′*^(*λ*) varies ∼ *λ*^4^/2, leading to:

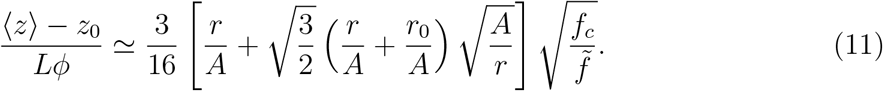

In this regime, the fluctuation dependent correction varies with force as 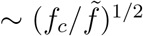, which is the same behavior as *z*_0_*/L* in 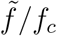; the fluctuation correction can be interpreted as changing the polymer persistence length *A*. The bulkier is the polymer (depending on the fraction *r*_0_*/A*) the greater is the contribution of fluctuations. Notably, as a function of *r/A*, the function has a minimum for crowders of radius *r* ≈ *r*_0_, and increases for larger *r/A*.

Requiring a smooth match of the two asymptotic behaviors of the function in *r/A* implies that, at fixed 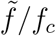, the full form of (⟨*z*⟩ − *z*_0_)/(*Lϕ*) from Eq. 10 has a maximum for the crowders of size close to the force-dependent correlation length *r* ≈ *ξ*. In fact, this is true only for slender enough polymers (when 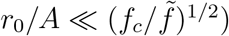, as can be seen in Fig. 2D. This also implies that, at high enough forces, the swelling effect of fluctuations generated by small crowders is always larger to the one by larger crowders, although it is so small in absolute terms that it cannot be seen in Fig. 2C.

When we consider the combined effect of the zeroth-order collapse and the fluctuation dependent swelling, strikingly, we predict that overall, crowders can *extend* the chain, increasing extension above the values for *ϕ* = 0, if the crowders are large enough such that *f*^∗^ < *f*_*c*_, and small enough such that the effect of fluctuations is significant (Fig. 2C, brown line), or for very high forces. This unexpected result belies the simple notion that neutral (purely steric) crowders can only compact a polymer. Experiments using Förster resonance energy transfer (FRET) experiments on protein disordered domains indicate that the same crowders can induce either increased folding or unfolding depending on the tested domain [21]. It is possible that these experimental results are due to physical effects discussed here, with compaction or decompaction dependent on the the size of crowders relative to the polymer bending stiffness, as well as the force applied to the chain. At present quantitative comparison to that experimental data is not possible, since the theory of this paper is valid only in the large pulling force regime (for *f > f*_*c*_) while solution FRET experiments are performed on unstretched polymers.

In this paper we have developed a theory of effects of steric crowders of radius *r* on the force-extension response of a strongly stretched DNA or other semiflexible polymer, including fluctuation effects through first order in crowder fraction *ϕ*. Our results predict *in vitro* experiments which could be used as a sensitive probe of crowder effects relevant to compaction of chromosomes or other polymer structures *in vivo*.

As a function of crowder size and volume fraction *ϕ* we find that there is an additive shift of net force on the polymer by *f*^∗^ ≈ *k*_*B*_*Tϕ/r* which can drive a collapse against external force, plus changes in the high-force response including a regime where the apparent persistence length is changed. These effects will be important to experimentalists since the force exerted by the apparatus will no longer be the net tension stretching the polymer once crowders are present. For example, at fixed external force *f*, addition of crowders can lead to a shift down of the apparent force and enhanced transverse fluctuations in experiments that calibrate force via fluctuations (e.g., magnetic tweezers [17, 22]), or alternately a shift up in force in cantilever-type experiments that essentially maintain constant extension (e.g., optical tweezers, atomic force microscopes). For optical tweezers, attention should also be paid to constant-force system dynamics, as well as to laser heating and trapping effects on the crowders themselves. Exploration of the different regimes of crowder size and force relative to the force characterizing the boundary between different fluctuation effect scaling, *f*_*λ*_ ≈ *k*_*B*_*TA/r*^2^, can modulate these basic effects and in some cases reverse them.

Our analysis uses approximations that need to be considered carefully when comparing with experiments. We use the limit of small crowder volume fraction when computing osmotic pressure, and the perturbation expansion for the WLC partition function, using the strong stretching regime ⟨*z*⟩ *> L*/2 where bending fluctuations are relatively small. Given that cellular and experimental levels of crowders are in the volume fraction range from 10 to 30 %, it would be of interest to include nonideal gas (*e*.*g*., using the Carnahan-Starling equation approach [23]) effects to the theory. Also, extension of the theory to describe the presumably crowding-collapsed globule that forms beyond the instability that we predict, would be of interest since crowding-induced polymer collapse has been experimentally observed.

Even in its current simple form, considering the broad range of forces over which we predict quantitative and qualitative effects, our calculations should be useful in understanding *in vivo* compaction of bacterial nucleoids and other intracellular structures. For instance, while it is accepted that RNA and DNA polymerases, structural maintenance of chromosomes complexes, and other molecular motors exerting forces on DNA can change chromosome condensation due to motor effects [24], our theory demonstrates that effective forces are sensed by a polymer, in response to varied crowder size and volume fraction.

We also observe condensation effects without any globular phase for physiological tensions, which raises the question of whether the bacterial nucleoid is condensed on its own, or rather is decondensed with size dependent on activities and local environment. These two options are not necessarily mutually exclusive, since it is possible to have a polymer coexist in an activity induced decondensed phase as well as more compacted regions in a different genomic-scale compartment, consistent with results of Hi-C and ATACseq genomic experiments on human cell lines [25, 26].

Consideration of *in vivo* conditions is complicated by the fact that the cellular environment is not simply a solution of spherical monodisperse crowders. More specifically the intracellular environment possesses *(i)* a high degree of polydispersity, *(ii)* heterogeneity of genomic domains in terms of binding proteins and their post-translational modifications, and *(iii)* presence of molecular activities in the cytoplasm. These factors are an objective for further development of the theory of the present paper to understand experimental results [14].

## ACKNOWLEDGMENTS

The authors acknowledge helpful discussions with Quentin Chaboche, Marco Cosentino-Lagomarsino, Sumitabha Brahmachari, Jean-François Joanny, Françoise Brochard-Wyart and Antoine Coulon. VFS acknowledges Laura Finzi, David Dunlap, Jin Qian, and Narendar Kolimi for sharing preliminary optical trap force-extension data. This work was supported by the Centre National de la Recherche Scientifique (CNRS), France (VFS), and by the NIH (US) through grants R01-GM105847 (JFM). JFM and VFS also acknowledge travel support from the CNRS “Physics Meets Genomes” Conference and Summer School in Cargese (France) where this work was initiated.

## APPENDIX A

The effective energy in Eq. 1 can be derived from a Hamiltonian that considers the configuration of the crowders in space. We add an interaction term between each crowder of position **x**_*i*_ and the polymer chain to a classical WLC energy for the polymer chain. Individual crowders are indexed by *i* spanning from 1 to *N*, such that the concentration of crowders is *c* = *N/V*, where *V* is the full experimental volume. This results in:

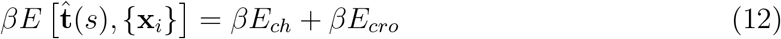

with

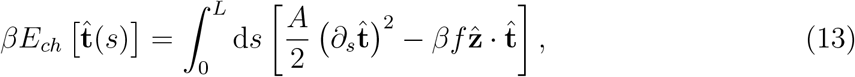

and

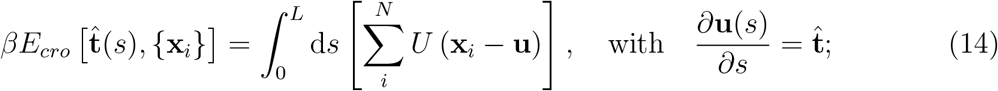

this is valid for low crowder densities (low enough *ϕ*), since it ignores two-body interactions between crowders, as well as higher orders; further, this is valid for overdamped conditions, when the inertial mass of the crowders and the chain are both negligible. Given monodisperse crowders of radius *d*, we set the steric interaction potential as:

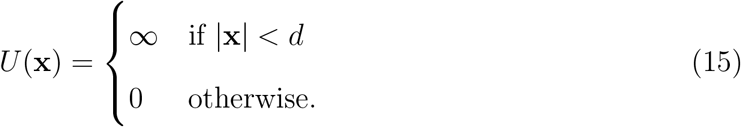

we can then derive the corresponding partition function from the Gibbs distribution, that is a path integral over all worm like chain configurations, integrated over all the positions of the crowders over the volume *V* :

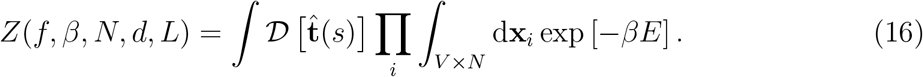

We then integrate the partition function over the crowder degrees of freedom {**x**_*i*_}. This allows calculation of the reduction in configuration space available due to the interaction with the polymer chain, i.e., the strength of depletion interactions. Considering the shape of the potential *U* (**x**), the problem reduces to calculating the reduction of volume accessible to the crowders 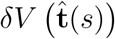 . In the specifics,

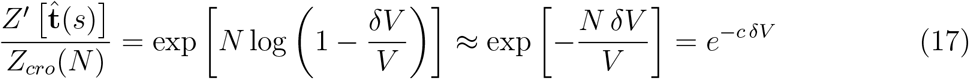

that is the van’t Hoff entropy approximation [27], valid for *δV* ≪ *V* . More realistic forms of *U* (**x**) would lead to a similar result. Of important note, this approximation not only simplifies the calculation, but also assures that *c* is not affected by variations in volume *δV* . Substituting, we obtain:

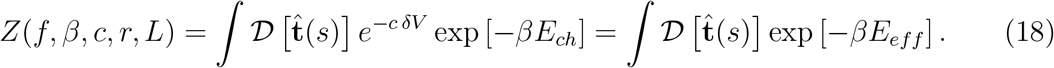

as defined in Eq. 1.

